# DNB-Based On-Chip Motif Finding (DocMF): a High-Throughput Method to Profile Different Types of Protein-DNA Interactions

**DOI:** 10.1101/827428

**Authors:** Zhuokun Li, Xiaojue Wang, Dongyang Xu, Dengwei Zhang, Dan Wang, Xuechen Dai, Qi Wang, Zhou Li, Ying Gu, Wenjie Ouyang, Shuchang Zhao, Baoqian Huang, Jian Gong, Jing Zhao, Ao Chen, Yue Shen, Yuliang Dong, Wenwei Zhang, Xun Xu, Chongjun Xu, Yuan Jiang

**Author notes:** To whom correspondence should be addressed., Correspondence may also be addressed to Chongjun Xu., Correspondence may also be addressed to Xun Xu. Joint Authors (Zhuokun Li, Xiaojue Wang, Dongyang Xu, Dengwei Zhang).

## Abstract

Here we report a highly sensitive DNB-based on-chip Motif Finding (DocMF) system that utilizes high throughput next-generation-sequencing (NGS) chips to profile protein binding or cleaving activity. Using DocMF, we successfully identified a variety of endonuclease recognition sites and the protospacer-adjacent-motif (PAM) sequences of different CRISPR systems. Our DocMF platform can simultaneously screen both 5’ and 3’ PAM regions with high coverage using the same NGS library/chip. For the well-studied SpCas9, our DocMF platform identified a small proportion of noncanonical 5’-NAG-3’ (∼5%) and 5’-NGA-3’ (∼1.6%), in addition to its common PAMs, 5’-NGG-3’ (∼89.9%). We also used the DocMF to assay two uncharacterized Cas endonucleases, VeCas9 and BvCpf1. VeCas9 PAMs were not detected by the conventional PAM depletion method. However, DocMF discovered that both VeCas9 and BvCpf1 required broader and more complicated PAM sequences for target recognition. VeCas9 preferred the R-rich motifs, whereas BvCpf1 used the T-rich PAMs. Moreover, after slightly changing the experimental protocol, we observed that dCas9, a DNA-binding protein lacking endonuclease activity, preferably binded to the previously reported PAMs 5’-NGG-3’. In summary, our studies demonstrate that DocMF is the first tool with the capacity to exhaustively assay both the binding and the cutting properties of different DNA-binding proteins.

## INTRODUCTION

DNA and proteins are the two most important biological macromolecules in organisms, and their interactions play crucial roles in many living cell activities, such as gene expression, DNA replication, viral infection, etc. DNA-protein interactions are necessary to translate the encoded genetic information for use by the cells. A major function of protein-DNA interactions is in the regulation of DNA architecture. These associations between DNA and protein primarily involve binding interactions. Proteins can interact with DNA in either the major or minor groove and in a sequence-specific or secondary structure-dependent manner, often inducing large structural changes in DNA. In both prokaryotes and eukaryotes, some proteins such as nucleases bind and subsequently cleave scissile phosphodiester bonds in nucleic acids, which is essential for biological processes like DNA repair or cell defence (1).

Current methods for studying DNA-protein interactions include electrophoretic mobility shift assays (EMSAs), chromatin immunoprecipitation (ChIP), and DNase footprinting (2)(3)(4), but these methods can be laborious, are not high throughput, and can only be used in cases where the DNA remains intact after the interaction occurs. Protein-binding microarrays (PBMs), on which proteins bind to double-stranded oligonucleotides, have been used to study transcription-factor DNA binding site preferences (5). Although PBMs are considered high throughput, this technology is limited by the number of features that can be placed on an array. The complete catalog of 10-mers (106 features) is the current approximate limit for array technology. However, many DNA-binding proteins such as zinc-finger proteins have recognition sites longer than 10 bp. SELEX (6) can achieve higher throughput in combination with NGS techniques. Although SELEX is a very useful in vitro technique, it is biased toward high-affinity binding motifs and does not disclose the full spectrum of binding preferences (7)(8). SELEX also cannot be used for proteins with nuclease activity or in situations in which the DNA does not remain intact after interaction with proteins. The NGS Illumina chip, which contains billions of dsDNA features on its surface, has been used to quantitatively analyze RNA-protein interactions (9). Recently, Jung et al. (10) used a chip-hybridized association mapping platform (CHAMP) to study protein-DNA binding preferences.

Revolutionary next generation sequencing (NGS) technologies have remarkably decreased the cost of genome sequencing, and they provide several hundred millions or billions of short DNA sequences on the surface of a flow cell for the study of protein-DNA interactions. In addition to Illumina’s HiSeq, NextSeq, and NovaSeq platforms, BGI’s BGISEQ-500 and MGISEQ-2000 sequencing platforms have been extensively used in applications such as exon sequencing (11) single cell RNA sequencing (12) small non-coding RNA analysis (13), and non-invasive prenatal testing (NIPT) (14). The technology underlying BGISEQ-500 or MGISEQ-2000 combines DNA nanoball (DNB) nanoarrays (15) with polymerase-based stepwise sequencing (DNBSEQ^TM^).

DNB-based on-chip motif finding (DocMF) is similar to HT-SELEX or CHAMP, but unlike other methods for studying DNA protein interactions, DocMF can provide information about protein binding at high throughput scales and in situations involving DNA strand cleavage. DocMF utilizes the DNBSEQ^TM^ technology (15) and sequential imaging to detect cleavage/binding motifs involved in protein-DNA interactions (Figure 1A). From the sequence information and the fluorescent signal change of individual DNBs, we can find all sequences that interact with the protein and identify the specific motifs via bioinformatics analysis (Figure 1B). In this report, we successfully identified the recognition sites of different types of restriction endonucleases. We also detected the PAM sequences of SpCas9 (5’-NGG-3’, 5’-NAG-3’ and 5’-NGA-3’) (16) and two novel CRISPR endonucleases (VeCas9 5’-NNARR-3’ and BvCpf1 5’-TYTN-3’) using a universal DNB pool for these different Crispr-Cas systems. We also queried the DNA binding preferences of dCas9 (17), a mutant Cas9 lacking endonuclease activity, using a slightly modified DocMF workflow. Our identification of the dCas9 binding sites NGG agrees with previous reports (17). In conclusion, our platform provides a high-throughput method to interrogate a wide variety of protein-DNA interactions. The utilities of our DocMF platform can be extended to other applications, such as on-chip identification of off-target sites for a CRISPR-Cas system, single stranded DNA cleavage sites, or transcription factor binding motifs.

**Figure 1.**
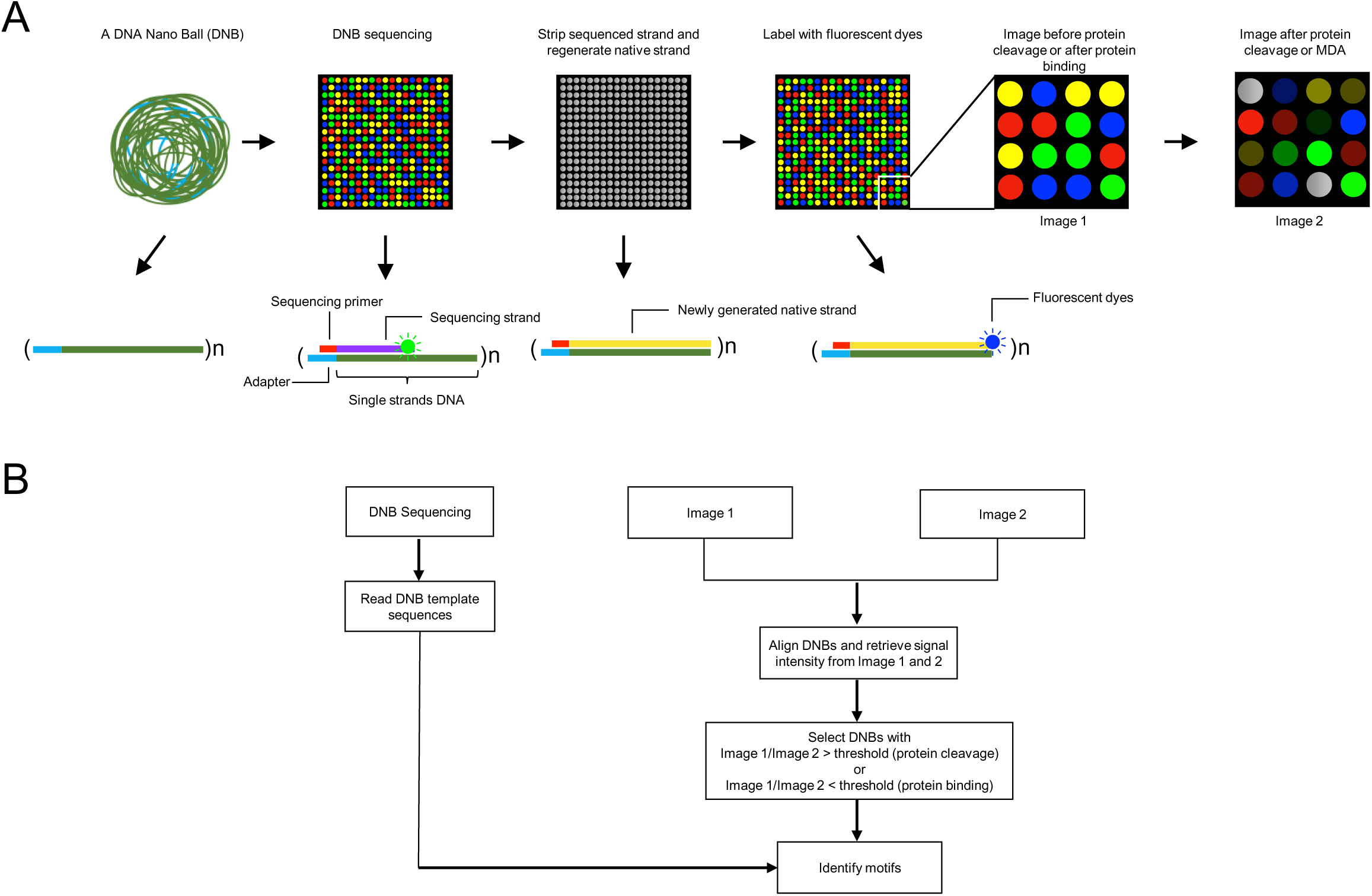
DocMF overview. (A) Biochemistry and illustration and (B) bioinformatics workflow.

## MATERIAL AND METHODS

### DNB library pool

For endonuclease assays, a synthetic oligo containing 50 random bases flanked by DNBSEQTM adapter sequences was purchased from Sangon Biotech Co. Next, 2 ng of the oligo was PCR amplified using 9 cycles of the PCR step in the MGIEasy Universal DNA Library prep set (MGI Tech Co., Ltd.). The PCR product was purified by bead purification according to kit instructions and quantified using the Qubit ™dsDNA High Sensitivity Assay kit (Thermo Fisher Scientific).

For PAM identification experiments and dCas9 binding experiments, the sequences of synthetic oligos used for library preparation were summarized in Supplementary Table 1 and all oligos were provided by China National Gene Bank. The double-stranded DNA used to make single strand circles was prepared by PCR amplification using the MGIEasy Universal DNA Library prep set (MGI Tech Co., Ltd.). In this PCR reaction, PAM_oligo_2-1 and PAM_oligo_2-2 were mixed and incubated at 95°C for 3 minutes and at 4°C for 10 minutes using a Bio-rad S1000TM thermocycler. This mixture served as PCR template, and PAM_oligo_1 and PAM_oligo_3 were used as primers. PCR was performed for 30 cycles according to the kit protocol. The PCR product was purified and quantified using the Qubit ™dsDNA High Sensitivity Assay kit.

Single-stranded circle (ssCir) preparation using 1 pmol PCR product from the previous step was performed according to the circularization step in the MGIEasy Universal DNA Library prep set.

Nuclease-free water (Ambion) was added to 6 ng ssCir DNA to achieve a 20-µl solution. Then 20 µl Make DNB buffer from the BGISEQ-500RS DNB Make Load Reagent Kit (MGI Tech Co., Ltd.) was added, and the mixture was incubated at 95°C for 1 minute, 65°C for 1 minute, and 40°C for 1 minute. Then, 40 µl make DNB enzyme mix V2.0 and 2 µl make DNB enzyme mix II V2.0 were added and incubated at 30°C for 20 minutes. The reaction was terminated by the addition of 20 µl DNB reaction stop buffer and immediate mixing.

### DocMF protocol

DNBs were loaded on the BGISEQ500 V3.1 chip by adding 30 µl BGISEQ500 DNB loader. Single-end sequencing runs for 55 bases were performed as instructed in the BGISEQ-500RS High-throughput Sequencing Set (SE100).

After sequencing, the labelled strand synthesized in sequencing was stripped off by 100% formamide (Sigma), a native complementary strand was synthesized on the BGIseq500 sequencer using dNTP mix II (MGI Tech Co., Ltd.), and dNTP mix I (MGI Tech Co., Ltd.) was used to add the final fluorescent dNTP.

DNB-protein interactions were assessed on chip using BGISEQ500 DNB loader. For PAM identification, the first images were acquired using Imaging reagent (MGI Tech Co., Ltd.) before treating DNBs with a protein of interest. For protein binding site identification, the first imaging was performed after treating DNBs with protein using the same protocol.

For PAM identification using DocMF, a second round of imaging was performed after protein-DNB interaction using the same imaging reagent. For protein binding motif identification, multiple displacement amplification (MDA) was performed after the first imaging, and then a second round of imaging was performed.

### Endonuclease restriction site characterization using DocMF

In this experiment, all restriction enzymes (EcoRI, Bpu10I, AgeI, NmeAIII, MluI, and BglI) were purchased from NEB.

The DNB library was prepared according to the description in the “DNB library pool” section. After native complementary strand synthesis, an image was captured.

Fifty units of the selected endonuclease was then pumped into the slide and incubated for 2 hours or overnight at the manufacturer recommended temperature for each endonuclease. The slide was washed with sequencing buffer, and the image after endonuclease digestion was captured.

### PAM identification using DocMF

In this experiment, the VeCas9 and BvCpf1 genes were subcloned into pET-28a vector to include an N-terminal His6 tag. Both genes were expressed in the E. coli BL21 (DE3) strain.

The templates for gRNA transcription were prepared by PCR. PCR templates, except for spCas9, are oligos ordered form IDT. For spCas9, plasmid PX458 was used as template. Primers used in these reactions were ordered from China National Gene Bank. The oligos used in these reactions are listed in Supplementary Table 1. PCR was performed using KAPAHiFi PCR HotStart Readymix (Roche) with an annealing temperature of 50° and an extension of 15 s for 30 cycles (melting temperature of one of two primers is 47°). The PCR products were purified using XP clean beads (Beckman Coulter) at a 2:1 ratio of beads to reaction volume. PCR products was quantified using the Qubit ™dsDNA High Sensitivity Assay kit (Thermo Fisher Scientific).

For guide RNA preparation, purified dsDNA from the previous step was incubated with T7 RNA polymerase overnight at 37°C using the MEGAscript T7 Transcription Kit (Thermo Fisher Scientific), and RNA was purified with MEGAclear Transcription Clean-up kit (Thermo Fisher Scientific).

crRNA was prepared by annealing two synthetic oligos (BvCpf1-crRNA-F and BvCpf1-crRNA-R) with complementary sequence ordered from China National Gene Bank. Obtained dsDNA was incubated with T7 RNA polymerase overnight at 37°C using the MEGAscript T7 Transcription Kit (Thermo Fisher Scientific). crRNAs were purified using RNAXP clean beads (Beckman Coulter) at a 2:1 ratio of beads to reaction volume with an additional 1.8x supplementation of isopropanol (Sigma).

All DNA oligo sequences used in this study are available in Supplementary Table 1.

All RNA sequences in this study are available in Supplementary Table 2.

The DNB-protein reaction mix was composed of 0.1 µM protein of interest, 3 µM corresponding guide RNA or crRNA, 1 µl RNase inhibitor (Epicentre), and nuclease-free water (Ambion) to a final volume of 300 µl. For Cpf1, the reaction buffer was NEB buffer 2.1. For Cas9, the buffer was NEB 3.1. The reaction mixture was loaded into the BGISEQ500 V3.1 chip by BGISEQ500 DNB loader and incubated for 4 hours at 37°C.

### dCas9 binding protocol

In this experiment, DNB-protein interactions were assessed on chip using BGISEQ500. For the experimental lane, the DNB-protein reaction mixture was composed of 0.1 µM dCas9 S. pyogenes (NEB), 3 µM corresponding guide RNA, 1 µl RNase inhibitor (Epicentre), 1X NEB buffer 3.1 (NEB), and nuclease-free water (Ambion) to a final volume of 300 µl. Before loading into the chip, the reaction mixture was incubated at room temperature for 15 minutes. Then the reaction mixture was loaded into the BGISEQ500 V3.1 chip by BGISEQ500 DNB loader and incubated for 4 hours at 37°C. For the control lane, 1X NEB buffer 3.1 (NEB) was used. A first imaging step was performed on the BGIseq500 sequencer (MGI) using Imaging reagent (MGI). After the first imaging, the MDA reaction was performed using 100 nM phi29 DNA Polymerase (NEB), 1x phi29 buffer (NEB), and 400 µM dNTP (NEB) with incubation at 30°C for 30 min. After the MDA reaction, a second imaging step was performed using the same protocol as the first imaging step.

### In vitro nuclease validation test

The cleavage substrate was amplified using overlap-PCR (PrimeSTAR GXL DNA Polymerase from Takara) from the random PAM plasmid library (see Supplementary Materials and Methods). A 300-bp fragment containing the spacer region was amplified by a pair of primers(M13R300 & PAMveriR, VeCas9-PAMveriR1-R10), and a 500-bp fragment containing part of the spacer region and the total specific PAM sequence was amplified using another pair of primers(M13F500 & VeCas9-PAMveriF1-18,BvCpf1-PAMveriF1-14, PAMveriF); the two fragments were subjected to overlap-PCR to obtain the substrate(M13F500 & M13R300). All designed oligos are shown in Supplementary Table 3.

VeCas9 and BvCpf1 were purified as described in Supplementary Materials and Methods. The required guide RNA sequence was obtained by small RNA sequencing. 100 ng of the cleavage substrate and a final concentration of 100 nM effector (VeCas9 or BvCpf1) and gRNA were used in a 20-μL reaction system. The reaction buffer system is Cas9 Nuclease Protein buffer (Abm). The reaction mixture was incubated at 25°C for 10 min before the addition of the cleavage substrate, and the cleavage reaction was conducted at 37°C for 1 hour. The reaction product (10 μL of total product) was detected using a 1% agarose TAE gel running at 150 V for 30 minutes.

### Processing DNB-based data

For each DNB, we obtained its read sequence and fluorescence intensity in Image 1 and Image 2 (Figure 1). We used the fold change of fluorescence intensity (FFI) to quantify the cleaving/binding effect for each DNB.

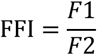

### Data analysis

#### Restriction endonuclease

To properly identify the exact length of restriction sites (RSs) for different restriction endonucleases, a “seed assembly” method was used. All 4-mer to 8-mer sequences were extracted from the 40-nt positive reads. Sequences with 1.5-fold or greater enrichment relative to the reference genome were regarded as seeds. Seeds with the same length were assembled with preference for the longest sequences. The length of the consensus sequences in all longest sequences was regarded as the predicted length of the enzyme recognition site.

After obtaining the length of the enzyme’s RS (L), we calculated the site rates of all possible L-mers using the following algorithm:

Suppose n is the initial total number of positive reads,

Step 1: Calculate the frequencies of all possible L-mers among positive reads via Equation 1 and only select the L-mer with the largest frequency. We designate this L-mer with the largest frequency as M. The site rate of M is determined and equals its frequency.

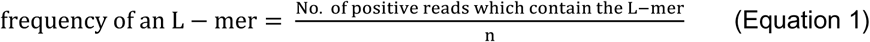

Step 2: Remove the reads which contain M from positive reads and repeat Step 1 until the site rates of all possible L-mers is obtained or there are no positive reads left.

#### Cas9, VeCas9, and BvCpf1: PAM identification and visualization

After selecting eligible reads with the correct protospacer, the 7-nt sequences at the 5’ and 3’ ends of the protospacer were extracted from these reads. Counting the total number for each unique 7-nt sequence, the relative read frequency was computed via Equations 2 and 3.

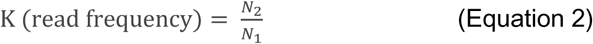

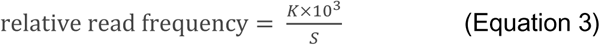

N1 is the total number of one particular 7-nt sequence for image 1, N2 is the total number of that 7-nt sequence for image 2, and S is the sum of whole K. By comparing the relative read frequencies at the 5’ and 3’ ends, we discovered that the read frequency of a partial 7-nt sequence at one end was much higher than its counterpart at the other end, indicating that the nuclease could recognize and cut these 7-nt sequences. Thus, the overall read frequency at the lower frequency end was regarded as background noise and an internal control. Because the read frequency of the control group, at the 5’ end or the 3’ end, showed a normal distribution, we applied the “three sigma rule” to define the cutoff shown in Equation 4.

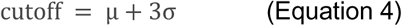

μ is the mean of the distribution of control, and σ is its standard deviation. A small portion of erroneous data could be excluded via this approach. The positive 7-nt sequences with higher read frequency were used to generate sequence logos by ggseqlogo (18). Similarly, Krona plots were plotted using these 7-nt sequences with t8heir read frequencies and further modified by Adobe Illustrator to produce a PAM wheel (19). To more accurately define the cutting efficiency of each 7-nt sequence, Fisher’s exact test was adopted as the main statistical method to prioritize the positive 7-nt sequence using the following formula (Equation 5):

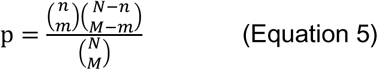

N represents the total number of image 1 for each 7-nt sequence, n represents the total number of image 2, M represents the number of unique 7-nt sequences in image 1, and m represents the corresponding number in image 2. Benjamini & Hochberg (BH) correction was used to control the false discovery rate as the multi-test adjustment method. All data were processed by Python and Excel.

For the frequency plot, each base frequency per site of 7-nt sequence in both image 1 and 2 was computed. Then, the base frequency of image 1 was subtracted from that of image 2 to discover whether the nuclease was functional.

#### dCas9

To analyze the data for dCas9, we used the “relative binding strength” (RBS) shown in the following equation to evaluate the binding strength for each 7-nt sequence.

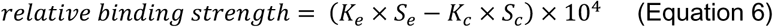

Ke is the read frequency for one particular 7-nt sequence in the experimental group, and Kc is its read frequency in the control group. Se is the sum of all Ke in the experimental group, and Sc is the sum of all Ke in the control group. The “three sigma rule” was also adopted to define the cutoff because the RBS at the 5’-end approximately fits a normal distribution. With this approach, we can extract the positive 7-nt sequences with a higher RBS. Python was used to split sequencing reads and extract 7-nt sequences, and data processing and plotting were completed using Excel and R, respectively.

## RESULTS

### Overview and optimization of DocMF system

The novel DocMF system measures protein:DNA interactions by examining the fluorescence signal change via on-chip sequential imaging before and after protein interaction with DNA targets (Figure 1). Hundreds of millions of DNA Nanoballs (DNBs) are first loaded onto the BGISEQ500 chips in a patterned array and sequenced at a fixed length using the DNBSeq^TM^ workflow (15). The DNBs contain random sequences to cover the full range of candidate protein binding sites. After obtaining the unique sequence information for each DNB, we reform single-stranded DNBs (ssDNBs) by stripping off the dye-labelled strand synthesized in sequencing. Subsequently, a native complementary strand is resynthesized to form dsDNA and end-labelled with fluorescent dyes. For DNA-cleaving proteins, A first image is acquired to record the location and the signal intensity of individual DNBs, i.e., reads. The protein of interest binds to its dsDNA targets and cleaves corresponding DNBs, leading to signal reduction or elimination during a second round of imaging (Figure 1A). Specific motifs can be identified from the sequences of selected DNBs with signal elimination or reduction greater than a threshold (Figure 1B) and verified in subsequent molecular assays. A slightly modified DocMF workflow is used to characterize protein:DNA binding preferences. In this protocol, DNBs are imaged after incubating end-labelled dsDNA with DNA binding proteins. In the following step, an additional polymerase reaction called multiple displacement amplification (MDA) is performed to replace the labelled strand. MDA leads to signal loss during the second imaging step as illustrated in Supplementary Figure 1. However, if a protein of interest binds to its DNA targets and inhibits MDA, the signal from the DNBs containing protein binding sites would remain unchanged or be less affected than that of the control lane, which does not include protein incubation but retains the other steps.

To ensure sequential imaging is feasible, it is crucial that the stripping step does not affect spatial information or damage the DNB structure. The BGISEQ500 chips used in this experiment are patterned arrays. Therefore, the sequential imaging does not affect the registration of DNB locations to the same extent that the CHAMP method using Miseq chips is affected (10). Additionally, we tested a variety of stripping buffers and found that the formamide buffer had the least impact on DNB integrity, only breaking the hydrogen bonds between dsDNA without affecting DNB stability or detaching DNBs from the surface. Supplementary Figure 2 shows that the sequencing quality scores, including Q30 (92.83 VS 90.32), Lag (0.15 VS 0.15) and RunOn (0.15 VS 0.12), remained unaffected after stripping with formamide buffer. In comparison, the stripping buffer with NaOH significantly decreased the Q30 from more than 90% to nearly 0 (data not shown).

After obtaining two images before and after protein:DNA interaction, we directly compared the raw signal intensity fold change of each DNB. If the protein cleaves DNA, the DNBs that have significant signal reduction can be retrieved (Figure 1B), and the corresponding sequences are analyzed for motif identification (Materials and Methods). To measure protein:DNA binding interactions, we obtained the binding sequence information from these DNBs with minimal signal fold change compared to the control.

### DocMF can characterize a broad range of restriction endonuclease recognition sites (RSs)

After the system was established, we tested six restriction endonucleases (EcoRI, Bpu10I, AgeI, NmeAIII, MluI, and BglI) with different RS features. The type II restriction enzymes cleave DNA adjacent to or within their recognition sites (20) which have been extensively studied. The selected enzymes contain RSs ranging from 6 bp to 11 bp and comprising normal palindromic sequences, non-palindromic sequences, and degenerate bases (Figure 2D). The DNB library contains a pool of synthetic random DNA fragments with a length of 50 nt. Forty of the 50 nucleotides of these random sequences were read using the BGISEQ-500RS High-throughput Sequencing Set (SE100). Images were taken before and after on-chip incubation of these enzymes for two hours or overnight. DNBs with fold change of fluorescence intensity (FFI) > 2 threshold (positive reads) were identified to screen for motifs (See Materials and Methods).

**Figure 2.**
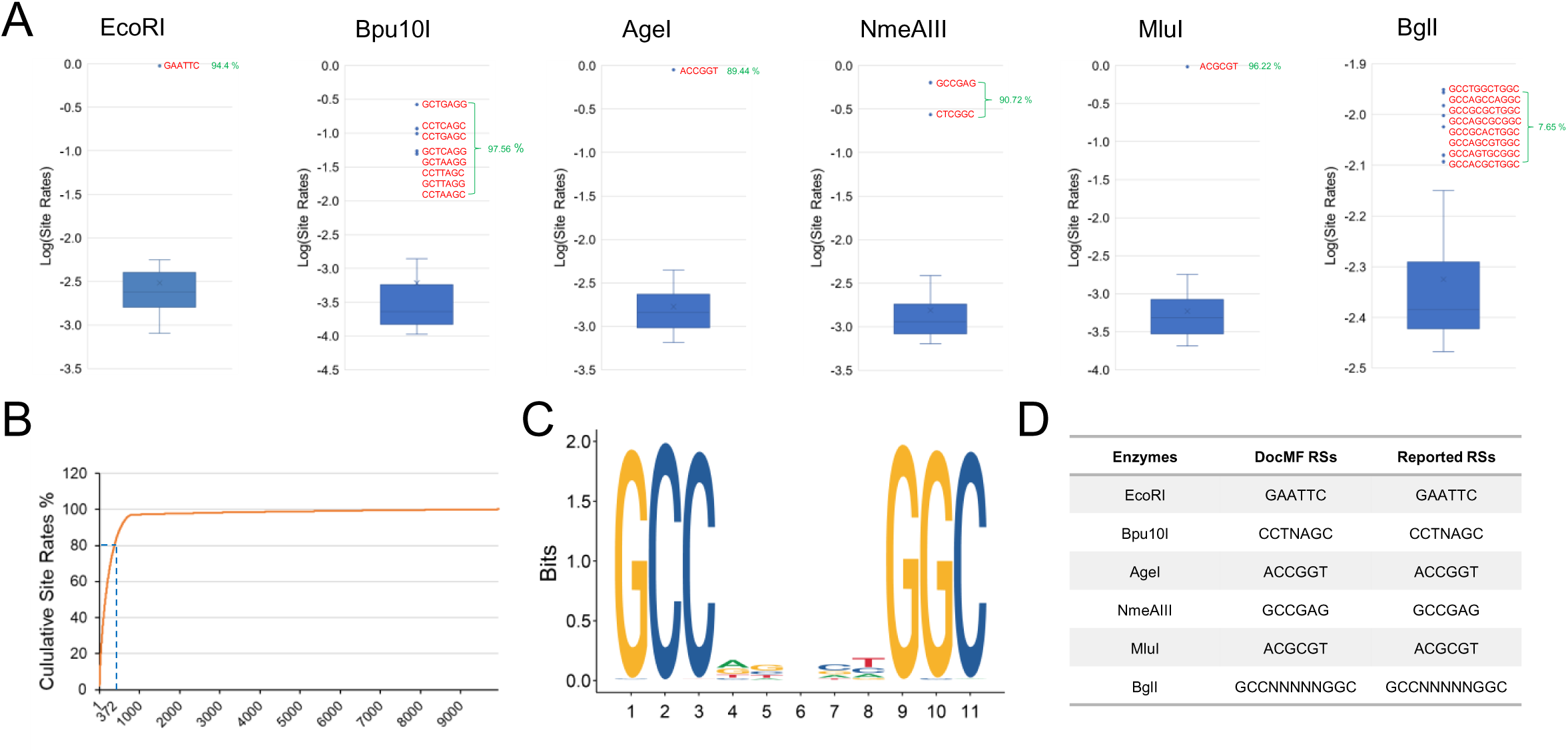
Restriction endonuclease cut site identification using DocMF. (A) Box plots for the motifs with the top 50 Log10(site rates). Outliers’ DNA sequences (orange) and the sum of outliers’ site rates (green) are shown. (B) Cumulative site rates for BglI. (C) A sequence logo representation of the 372 motifs for BglI. (D) Predicted DocMF RSs and reported RSs of the six restriction endonucleases.

The exact length of RSs (L) was obtained via a “seed assembly” method (Materials and Methods). We then calculated the site rates of all L-mers among positive reads (Materials and Methods). Using this method, for each of these six restriction endonucleases, we obtained the site rates of all L-mers (L is the predicted length of an RS) and drew a boxplot for the L-mers with the top 50 largest site rates (Figure 2A). The motifs (colored orange in Figure 2A) corresponding to the outliers in the boxplot, with the sum of site frequency of these outliers (colored green in Figure 2A) > 80%, were regarded as the DocMF predicted RSs. Thus, for EcoRI, Bpu10I, AgeI, NmeAIII, and MluI, we obtained their RSs from the outliers shown in Figure 2A, because the site rate sums of these outliers were all larger than 80%. For BglI, however, the site rate sum of the outliers was only 7.65%, which is too small to be predicted as RSs using DocMF. Thus, for BglI, we used the 11-mers (because 11 is the predicted length of the RS) with the top 372 largest site rates to predict the RSs; the site rate sum of these 372 motifs was larger than 80% (Figure 2B). A sequence (18) representation (Figure 2C) of these 372 sequences revealed that the RS for BglI was GCCNNNNNGGC, which agreed with the reported RS. These summarized results in Figure 2D demonstrated that our system and an optimized bioinformatics method could reliably identify the DNA recognition site for different types of restriction enzymes.

### DocMF can accurately identify the 5’-NGG-3’ PAM of SpCas9

CRISPR-Cas effectors are RNA-guided endonucleases that use a protospacer adjacent motif (PAM) as a DNA binding signal. The PAM is a short DNA sequence, normally less than 7 base pairs, that sits near the target DNA (termed protospacer) of the Crispr-Cas system (21)(22). One widely used PAM identification approach transforms plasmids carrying randomized PAM sequences into Escherichia coli in the presence or absence of the CRISPR-Cas locus. The frequency of a functional PAM sequence is significantly lower when the Cas protein is present (23).Thus, this PAM depletion assay (23) requires two sets of libraries for either 5’ or 3’ PAM identification and corresponding negative controls. The library size also needs to be large to cover most, if not all, PAM sequences. Additionally, the plasmid depletion assay (23) is time-consuming and low throughput. In contrast, the DocMF system can simultaneously screen both 5’ and 3’ sequences for PAMs in a single experiment, generating coverage that is multiple orders of magnitude greater than the traditional method. One of the two PAM regions that is not recognized by the protein is used as an internal negative control.

In a proof-of-concept study, we evaluated the accuracy of DocMF by assessing the PAM requirements of SpCas9, the most widely used CRISPR-Cas system, from Streptococcus pyogenes. SpCas9 cleaves the double-stranded DNA after binding to corresponding RNA, and this cleavage is reported from PAM depletion assays to be dependent on a 5’-NGG-3’ PAM sequence (24). The PAM DNB library used in DocMF is shown in Figure 3A. The synthetic oligo region contained a known 23-nt SpCas9 protospacer sequence (colored orange in Figure 3A) flanked by 5’ and 3’ PAM regions with 15 random nucleotides each (colored green in Figure 3A). The sequence information of both PAM regions was obtained by a single-end sequencing of 50 nt.

**Figure 3.**
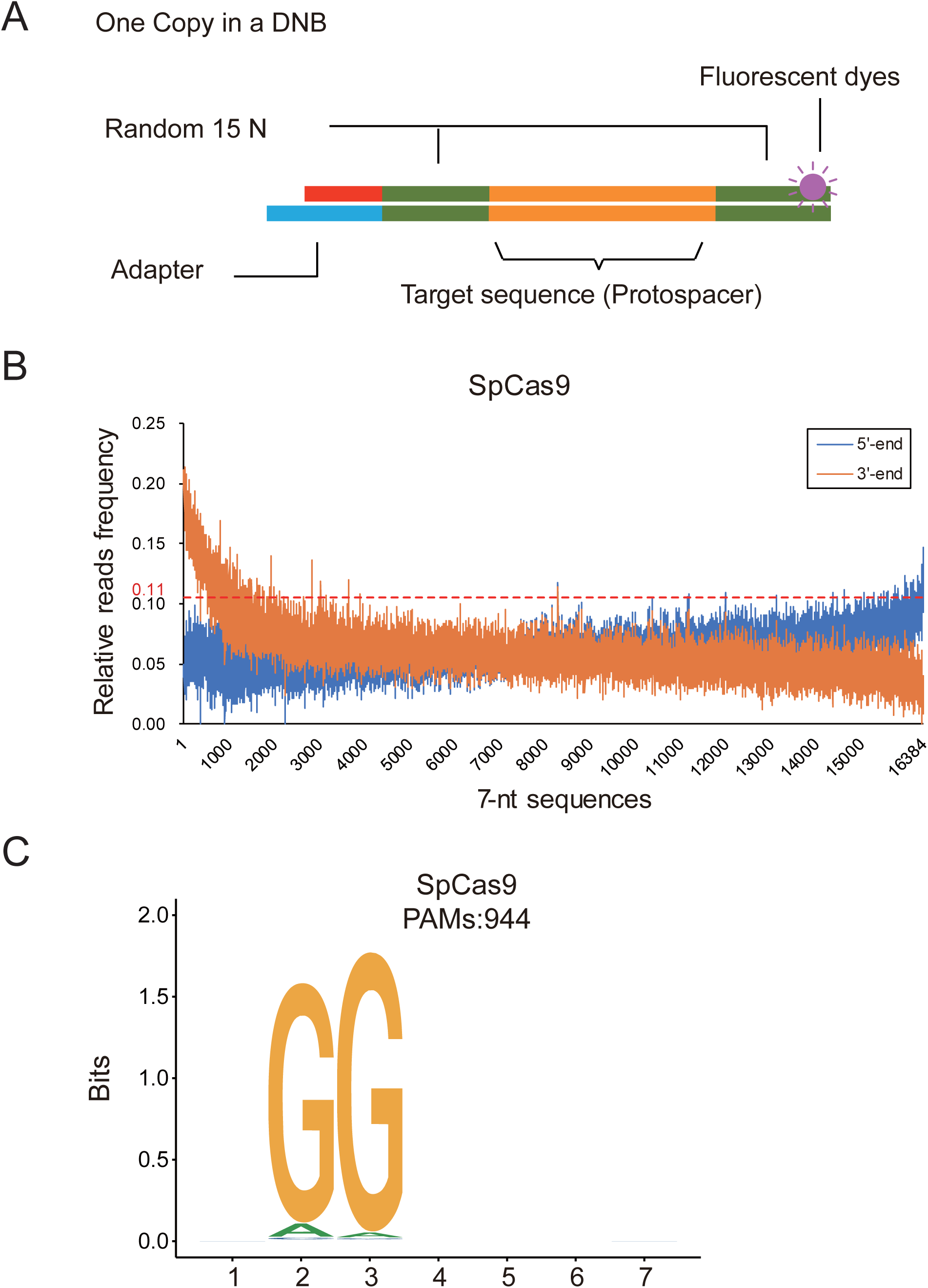
PAM identification for SpCas9 using DocMF. (A) PAM DNB library preparation illustration. The synthetic oligo region contains a known 25-nt SpCas9 protospacer sequence (orange) flanked by 5’ and 3’ PAM regions with 15 random nucleotides each (green). Hundreds of copies of each random-PAM-flanked protospacer are incorporated per DNB and only copy is demonstrated. (B) The relative read frequency at both the 5’-end and the 3’-end for SpCas9. The X-axis is all combinations of 7-nt sequences sorted by the difference between two ends in descending order. (C) PAM sequence for SpCas9.

The signal fold change was compared before and after SpCas9 on-chip incubation for 4 hours. Out of 494,866,059 reads (DNBs), we obtained 366,913 DNBs that exhibited a fold change greater than 3 and could potentially be cleaved by SpCas9. The 7-nt sequences at both 5’ and 3’PAM regions were retrieved from these DNBs for further analysis. The frequency of all 16,384 (4^7) PAM combinations for both 5’ and 3’PAMs was calculated and plotted against the individual sequence in Figure 3B. SpCas9 endonuclease was reported to only bind to the 3’ end of the target sequence. Therefore, 5’ randomized 7-nt sequences were used as the internal negative control. We applied the “three sigma rule” (25) to the 5’ sequences to define the cutoff for positive PAM signals. In other words, at the cutoff of 0.11, approximately 99.7% of data from the 5’PAM region fell into background noise. This statistical cutoff resulted in 944 3’PAM sequences that were preferably cut by SpCas9. A sequence logo (18) representation of the sequences revealed that SpCas9 preferred a 5’-NGG-3’ motif, although approximately 5.8% (55 out of 944) of 5’-NAG-3’ and 1.6% (15 out of 944) of 5’-NGA-3’ could also be recognized (Figure 3C), which is in line with previous findings (26)(27)(28).

### DocMF enables sensitive in vitro detection of PAMs in different Crispr-Cas systems

To further demonstrate the utility of DocMF in discovering PAM sequences, we extended our study to two previously uncharacterized Crispr-Cas systems, VeCas9 from Veillonella genus and BvCpf1 from Butyricimonas virosa (23). VeCas9 has a Cas9 effector protein of 1,064 amino acids (aa), and BvCpf1 protein is 1,245 aa in length. Both proteins were expressed and purified as described in Supplementary Materials and Methods. The crRNA and tracrRNA of VeCas9 were identified through small RNA sequencing, whereas the crRNA of BvCpf1 was predicted in silico based on the previously reported Cpf1 orthologs (Supplementary Figure 3). To interrogate the diversity of their PAM sequences, we conducted DocMF on VeCas9 and BvCpf1 using the same DNB PAM library (Figure 3A) used in the SpCas9 study. For VeCas9 experiments, we included three individual guide RNA designs, crRNA:tracrRNA, sgRNA-1 with SpCas9 structure, and a truncated sgRNA-2 (Supplementary Figure 3).

Prior to using DocMF, a PAM depletion assay (23) was first performed on VeCas9 for methodology comparison (Supplementary Materials and Methods). As shown in Supplementary Figure 4A-B, with 4.62 Gb of sequencing data, we observed 508 sequences with a threshold of 3 for Alicyclobacillus acidoterrestris C2c1 (AacC2c1), a positive control in the depletion assay, and correctly identified the reported PAM, 5’-TTN-3’ (23). However, with even more sequencing data (7.33 Gb) for VeCas9, 0 and 74 distinct sequences were found with thresholds of 3 and 0.8, respectively (Supplementary Figure 4C-D). The results for VeCas9 were quite similar to our negative control sets (data not shown), and thus we failed to detect correct PAM sequences for VeCas9 using the traditional depletion method.

Using DocMF, DNBs with signal fold change above threshold were selected for further analysis. In the read frequency plot (Figure 4A-B), the 5’PAM region of VeCas9 (with sgRNA-1) and the 3’PAM region of BvCpf1 showed no protein binding pattern, and their corresponding three standard deviations (0.09 for VeCas9 and 0.075 for BvCpf1) were used to set cutoff lines. As a result, 4,947 and 5,580 unique PAM sequences were determined to be cleaved by VeCas9 and BvCpf1, respectively. Both Crispr-Cas systems conveyed large PAM families as illustrated in consensus sequences and sequence logo, two common PAM reporting schemes (Figure 4C-D, E, G) (21)(22). The consensus sequences of VeCas9 were revealed as 5’-NNARRNN-3’, or NYARRMY for a even more dominant set of PAM sequences by frequency plot (Figure 4C), while sequence logo reported 5’-NNNRRNN-3’ PAM sequences (Figure 4E). VeCas9 with the other guide RNAs showed a similar pattern (Supplementary Figure 5). Over 99% of PAMs with the short sgRNA-2 were found with at least one of the other two RNAs, indicating the high reproducibility of this DocMF method. Slight difference in PAM of BvCpf1 was also observed between PAM reporting methods. Consensus sequence and sequence logo reported 5’-NNNTYTN-3’ (Figure 4D) or NNNNYYN (Figure 4G) respectively. However, these two reporting systems ignored the correlation among all seven positions and might introduce some incorrect active PAMs if randomly combining each position.

**Figure 4.**
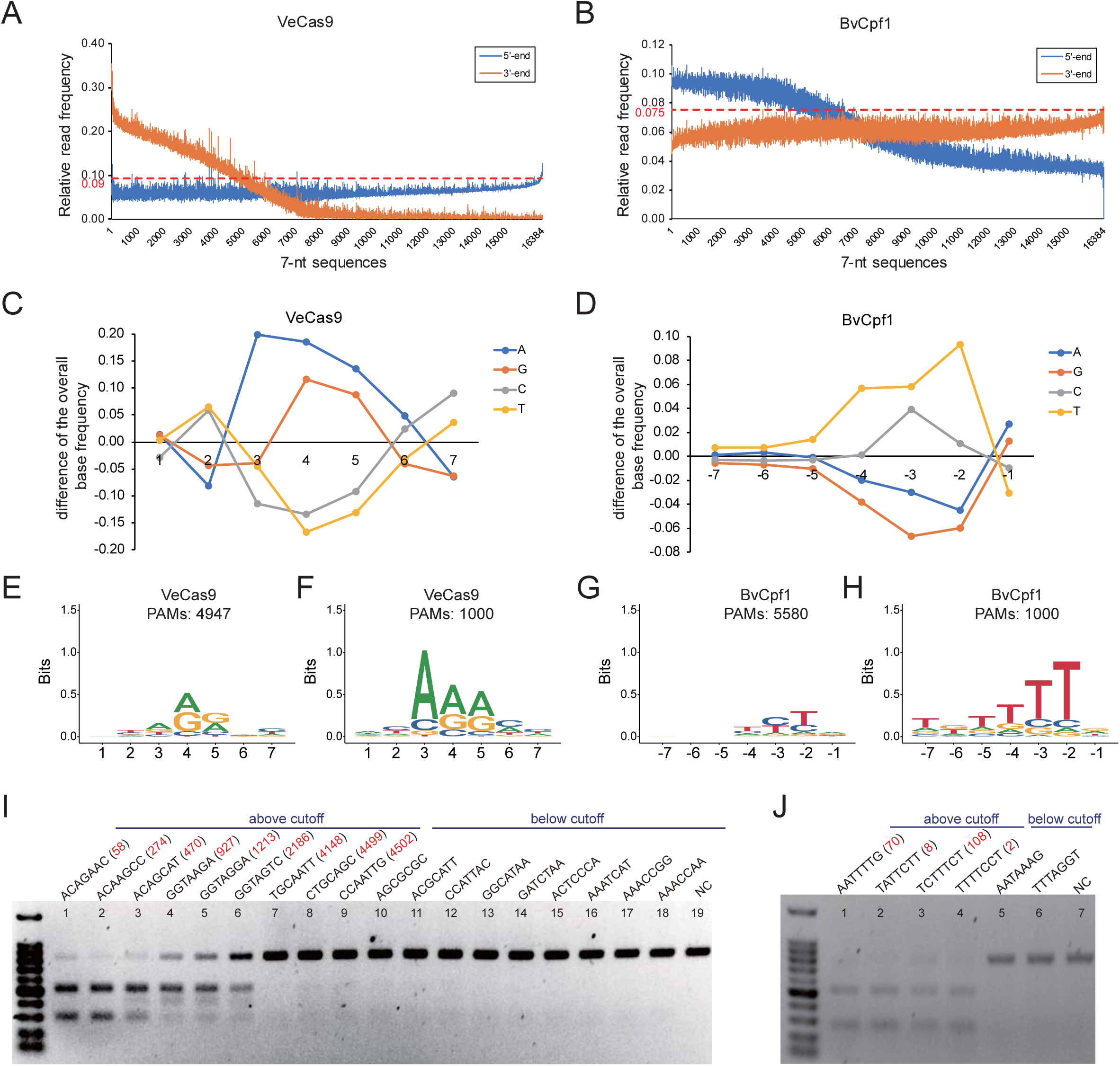
PAM identification in novel Crispr-Cas systems using DocMF. (A) The relative read frequency at both the 5’-end and the 3’-end for VeCas9. (B) The relative read frequency at both the 5’-end and the 3’-end for BvCpf1. Consensus PAM sequence by frequency plot with all detected 7-nt sequences for VeCas9 (C) and BvCpf1 (D). PAM sequence by sequence logo for VeCas9 generated by all detected 7-nt sequences (E) and by the top one thousand 7-nt sequences from FET analysis (F). PAM sequence by sequence logo for BvCpf1 generated by all detected 7-nt sequences (G) and by the top one thousand 7-nt sequences from FET analysis (H). (I) In vitro validation of VeCas9 PAM sequences. Nine 7-nt sequences each above/below the cutoff were selected. The FET ranking numbers are shown in red. NC means negative control. (J) In vitro validation of BvCpf1 PAM sequences. Five 7-nt sequences above the cutoff and two 7-nt sequences below the cutoff were selected.

To interrogate the relative activity of each PAM, two methods were applied, Fisher’s exact test (FET) and the PAM wheel. FET was introduced to sort the PAM sequences. FET is a widely used test to determine whether the difference between two groups is significant. Therefore, one particular PAM with a smaller p-value according to FET indicated that its relative read frequency, or cutting efficiency, was more significant compared with one with a larger p-value. After ranking the PAMs in order, we examined the consensus PAM sequences for the top 1,000 sequences (Figure 4 F, H). A slightly distinct PAM consensus sequence, 5’-NNARRNN-3’ for VeCas9 and 5’-NNNTTTN-3 for BvCpf1, was observed under this stringent selection criteria, which correlated better with the frequency plotting results (Figure 4C-D). To further validate the FET prediction, an in vitro nuclease assay was performed with randomly selected PAM sequences. PCR products containing individual PAMs and a common protospacer were incubated with either Cas9 or Cpf1 proteins at 37°C for one hour. Reactions with 50 ng of input were run on TAE gels, and the remaining input quantity was used to calculate cleavage efficiency. As demonstrated in Figure 4 (G-H), the PAMs with higher FET ranking numbers (in red) had less input remaining, indicating better cutting efficiency. The least ranked PAM gave minimal cutting, the product of which was almost not visible on agarose gels whose sensitivity are several orders of magnitudes lower than NGS. The consistency suggested that we could use our FET prediction to select the most active PAMs for in vivo gene editing.

A PAM wheel was also used to comprehensively understand the PAM sequences and their base dependence. Figure 5 A-B depicts the respective PAM wheels for VeCas9 and BvCpf1. For VeCas9, there is a strong base dependence between position 3 and 4. If position 3 had a base R (A or G), position 4 tended to have R (> 80%, Supplementary Figure 6) and a small but notable level of C (> 10%). If position 3 is Y (T or C), position 4 favoured R only (∼99%). T is the least favoured base at position 4 or 5, which agrees with the in vitro cutting results from Figure 5C (lane 9 and 10). The gel also demonstrated that NYARRMY (the consensus PAM based on the most dominant consensus sequence, lanes 1 in Figure 4C), NNARRNN (lanes 2-4) and ACAAGCC (58^th^ ranked sequence as positive control, lane 11) was cut more efficiently than NNCRRNN (lane 5), NNTRRNN (lane 6), or NNGRRNN (lane 7), which explains why A composed 47% at position 3 while the other three bases were each between 16% to 19% (Figure 5A, Supplementary Figure 6). For the BvCpf1 PAM wheel shown in Figure 5B, we discovered that position -4 was random when both positions -2 and -3 were Y (T/C). So PAM NNNNYYN generated more cutting products than NNNNYRN in Figure 5D (lane 1 and 2). However, position -4 tended to be T if one of the -2 or -3 positions was not Y. As a result, we observed slightly more cutting with T than V at position -4, when position -2 and -3 are either RY or YR (Figure 5D lane 7-10). R at position -2 also dictated that position -3 will be Y (100%, Supplementary Figure 6). As shown in Figure 5D, BvCpf1 system demonstrated clear cutting on 5’-NNNTTTN-3’ or NNNTYTN (Figure 5D). Our data suggested that both VeCas9 and BvCpf1 had a complicated set of PAM sequences which were comprehensively captured by DocMF.

**Figure 5.**
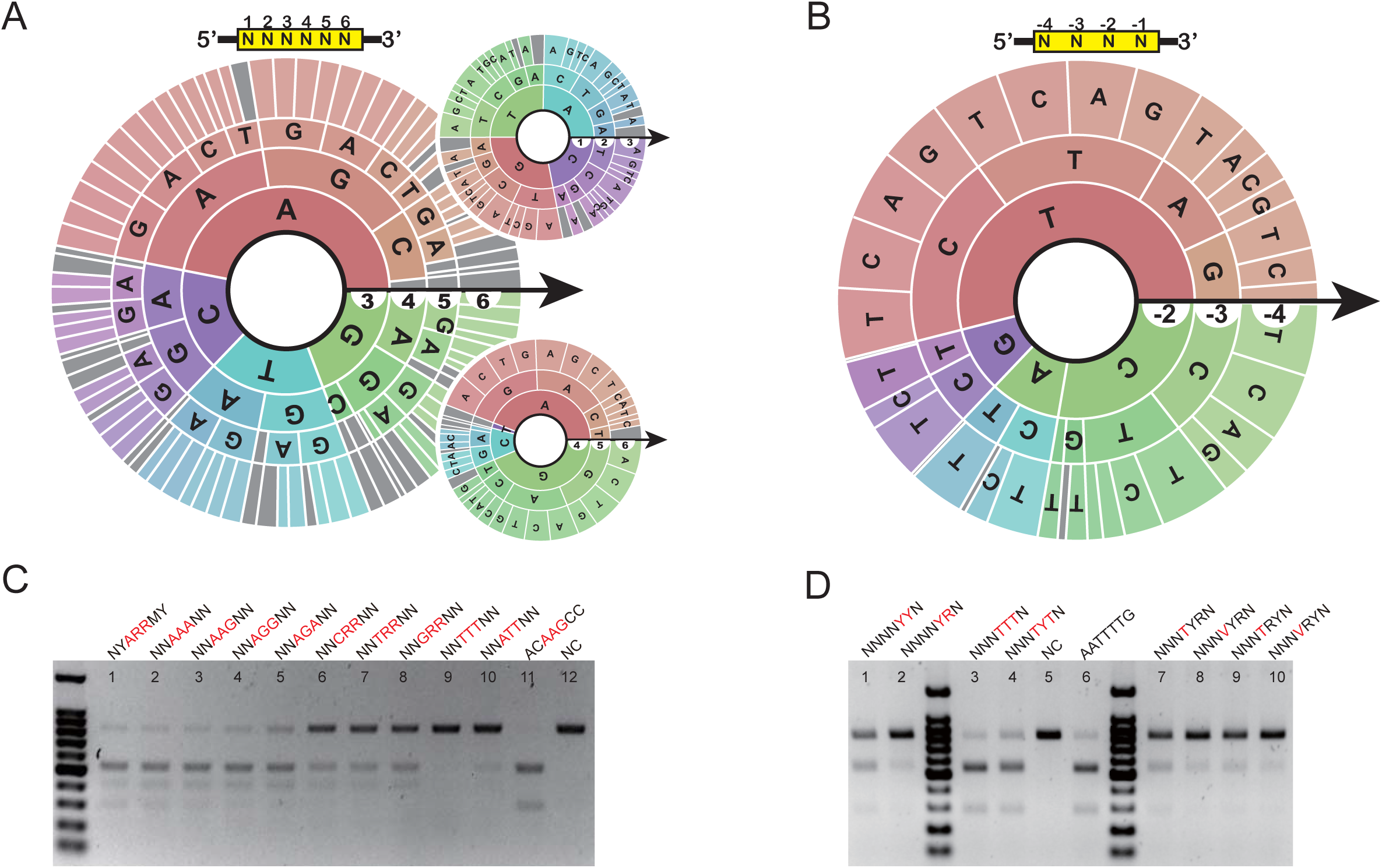
PAM wheel results. (A) PAM wheel for VeCas9. The upper yellow box gives an indication about each position of the PAM sequence, and the arrow illustrates the orientation of each base. The area of a sector of the ring for one base at one particular position represents its frequency at this position. (B) PAM wheel for BvCpf1. (C) In vitro validation of VeCas9 PAM wheel results. NYARRMY is the consensus sequence based on positional frequency, while ACAAGCC is 58^th^ FET ranked sequence included as a positive control. (D) In vitro validation of BvCpf1 PAM wheel results. NNNTTTN or NNNTYTN is the consensus sequence based on positional frequency, while AATTTTG is 70^th^ FET ranked sequence included as positive control. NC (Negative control): positive PAMs incubated with corresponding protein but without any sgRNA.

### DocMF can accurately identify protein binding sites

Protein-DNA interactions have been characterized in many high-throughput platforms including microarrays, HT-SELEX, and CHAMP (6)(10). We modified the DocMF workflow mentioned above to detect protein-DNA binding motifs. The steps remained unchanged until the natural complementary strand was resynthesized to form 50-bp dsDNA and end-labelled with fluorescent dyes. After binding the protein of interest to its dsDNA targets and washing off any excess, we acquired the first images to record signal intensity. After this first imaging, an on-chip incubation with MDA reaction buffer, dNTPs, and a polymerase with strong strand displacement was performed at 30°C for 30 min to synthesize a second complementary strand using the ssDNB as template (Supplementary Figure 1). Consequently, the original fluorescent strand would be replaced and displaced from DNBs, leading to signal drop when there was no protein binding to prevent MDA. To test this idea, we used a well-studied protein, dCas9, and removed its endonuclease activity through point mutations in its endonuclease domains HNH and RucV (16). The point mutations D10A and H840A changed two important residues for endonuclease activity, which ultimately results in its deactivation. Although dCas9 lacks endonuclease activity, it remains capable of binding to its guide RNA and the DNA strand that is being targeted because the binding is mediated through its REC1, BH, and PI domains (29). Moreover, dCas9 has previously been shown not to bind to its target sequence when there is no PAM (NGG) present (30). Unlike the studies above, the reads with fluorescence fold change below the threshold were considered positive, indicating the DNBs that dCas9 could recognize and bind. Additionally, we included a negative control lane without dCas9 incubation in the same process, since BGISEQ-500 has two lanes on a single chip. Approximately 95% of a total of 253M reads from the negative control lane lost half of the signal intensity (data not shown). We chose a signal change at 0.5 as threshold (Image 2/Image 1) and retrieved 14,371,289 out of 335,497,075 and 15,647,574 out of 337,529,837 reads from experimental and control lanes, respectively. We introduced a reliable “relative binding strength” concept to evaluate the binding strength for each 7-nt sequence. Similarly, the data at the 5’-end fitting a normal distribution were regarded as background noise, and the “three sigma rule” was adopted to define the cutoff at 0.135. After deducting the noises, we observed the NGG sequence was essential for dCas9’s binding (Figure 6), consistent with previous findings. This suggests that with the modified DocMF workflows can be harnessed as a general tool for identifying DNA binding motifs.

**Figure 6.**
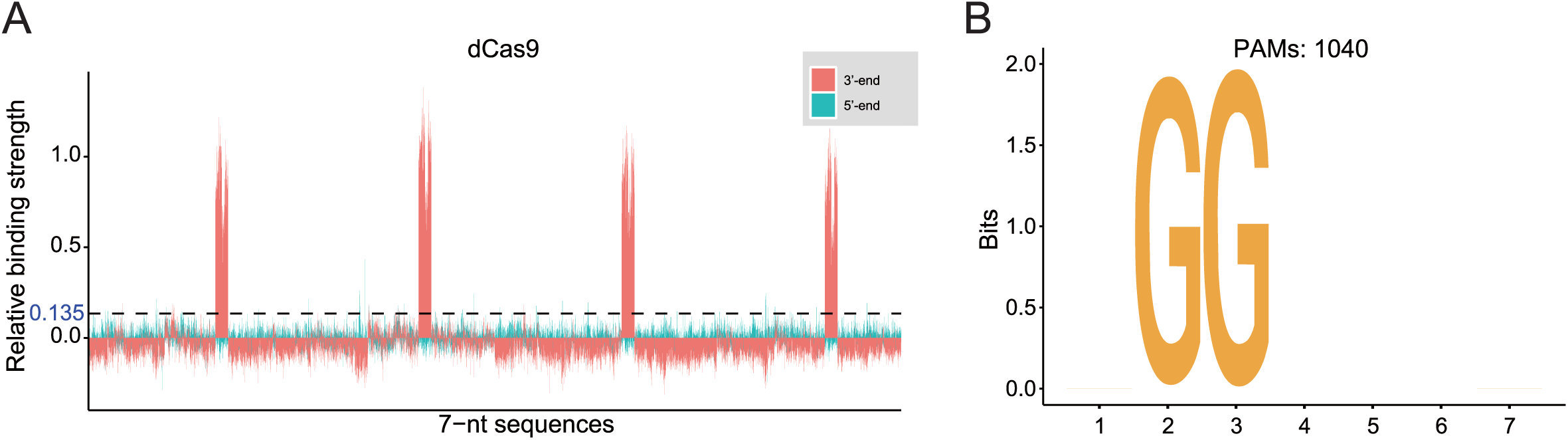
DocMF used to identify protein binding site of dCas9. (A) The relative binding strength at both the 5’-end and the 3’-end for dCas9. The X-axis is all combinations of 7-nt sequences and is automatically sorted by letter using Excel. (B) Sequence logo for dCas9 was generated by all detected 7-nt sequences based on those with the highest relative binding strength.

## DISCUSSION

Similar to CHAMP (chip-hybridized association mapping Platform (10), DocMF utilizes next-generation-sequencing chips to decipher protein-DNA interactions in a high-throughput manner. However, the two systems differ in many aspects. First, CHAMP needs to tag proteins with epitopes and uses fluorescent antibodies against the epitope to label the proteins on chip. In contrast, DocMF directly incubates proteins with dye-labelled target DNAs, enabling a simpler protocol and a cleaner result without the concern of non-specific noise from surface immunostaining. Secondly, CHAMP uses a random clustering chip and therefore needs a fluorescent alignment marker for cluster localization information. In contrast, DocMF is performed on chips with patterned arrays followed by a straightforward sequential imaging workflow to provide protein-DNA binding information. This approach significantly simplifies and expedites the on-chip biochemistry and the downstream bioinformatic analysis. Third, like many other high throughput technologies such as protein-binding microarrays and SELEX (6), CHAMP can only examine protein-DNA binding intensity. CHAMP researchers include ATP inhibitors to prevent Cas3 from digesting DNA clusters when assaying for Cas3 recruitment to the DNA-bound Cascade complex (10). However, DocMF is designed to explore various types of protein-DNA interactions, including both binding and cutting.

In this study, one important application of DocMF was to quickly and accurately identify the PAMs of any Crispr-Cas system (Figure 3-4). The Crispr-Cas system was initially identified in the prokaryotic immune system and was quickly adopted as a reliable genome engineering tool (24)(31). The PAM sequence is adjacent to the target site and is essential for Cas endonuclease specificity. Different Cas proteins bind to a variety of PAM sequences and exhibit different off-target rates of cleavage (23). To increase the number of potential genome editing sites, we are in urgent need of new Cas proteins that recognize PAM sequences beyond the commonly used 5’-NGG-3’ site for SpCas9 (16). DocMF could be a very useful tool for characterizing the novel Cas protein PAM sequences with the following advantages: 1) a universal system for different Crispr-Cas systems, with the same DNB pools containing a common 25-nt protospacer flanked by two 15-bp randomized sequences (5’ and 3’PAM regions), as demonstrated in this study for SpCas9, VeCas9 and BvCpf1 proteins; 2) high sensitivity, offering average of 20,000x-30,000x per unique sequence for a 7-bp PAM (400-500M reads for a total of 16,384 sequences) from a single BGISEQ-500 lane, compared to approximately 20x coverage from the E.coli completion assay; 3) inclusion of an internal negative control, i.e., the 15-mer that is not bound by Cas proteins, to better define true positives; and 4) high accuracy with validated cutting efficiency by in vitro assays. Using the DocMF system, we found that VeCas9, a new ortholog of Cas9 found in Veillonella sp., recognizes 5’-NNNRR-3’, especially 5’-NNARR-3’. The diverse PAM sequences of VeCas9 could potentially be advantageous in gene editing, especially when no suitable SpCas9 PAM is present. Additionally, with a size of 1,064 amino acid residues, VeCas9 can be easily packaged into adeno-associated virus (AAV), assisting better AAV delivery. The same DocMF system was applied to understand a Cpf1 ortholog, BvCpf1. The PAMs of BvCpf1 are T-rich sequences, but they are on the 5’ side of the common 25-nt target sequence. Cpf1 has recently emerged as another powerful tool for gene editing with features distinct from Cas9, such as a requirement for T-rich 5’ PAMs, a single guide RNA, and the production of a staggered DNA double-stranded break (31). Our characterization of VeCas9 and BvCpf1 helps to expand the existing CRISPR toolbox and provide more candidates for genome engineering. Moreover, diverse PAM sequences with a full spectrum of cutting efficiency can be obtained from DocMF. This remarkably sensitive sorting from our FET analysis can not only help researchers to identify the strongest cutting sites, but also predict potential off targets for their in vivo experiments.

Next, we performed a proof-of-concept study by assaying protein-DNA binding affinity using DocMF. dCas9, like many transcription factors (TFs), binds to DNA in a sequence-specific manner. In the modified DocMF workflow, an enzymatic reaction called MDA is added between the two imaging steps. MDA displaces the dye-labelled strand and therefore causes signal loss. However, if there is any dCas9 associated with the fluorescent strand, MDA mediated by phi29 polymerase will stop at the protein-DNA binding sites, leading to no or minimal signal change. In this experiment, we also ran a negative control experiment without dCas9 incubation on a separate lane. After removing the false positive sequences, we discovered that dCas9 exclusively bound to a motif of NGG. The NAG preference was not present for dCas9, indicating that the point mutations in the endonuclease domains, although outside of PAM-interacting (PI) domains, might also change the PAM specificity.

In the nucleus, TFs and their cofactors normally form multi-protein complexes and bind to chromatin through DNA binding motifs to mediate gene expression. The ease of adding other proteins through the fluidic system of the NGS sequencers also provides the potential to use DocMF to examine DNA interactions with protein complex. The only caveat of this protocol is that if the protein-DNA binding affinity is too weak to block MDA, we must first cross-link the protein and DNA using a simple formaldehyde fixation before MDA. The dissociation constant (KD) between dCas9 and the entire DNA substrate is estimated to be 2.70 nM (32). Therefore, TF-DNA interactions with KDs in the nM and pM range can be directly assayed with the DocMF protocols provided in this study.

In summary, DocMF, to our knowledge, is the first high-throughput platform that can characterize motifs for both DNA binding and cleaving proteins. DocMF offers high levels of accuracy and sensitivity in motif identification. In addition to the restriction site identification, PAM characterization, and DNA binding motif examination demonstrated in this study, the utilities of our DocMF platform can also be extended to predict other aspects of protein-DNA interactions, such as on-chip identification of off-target sites for any CRISPR-Cas system, single-stranded DNA cleavage sites, or binding motifs of protein complex-like transcription factors. This workflow can also be easily adopted by Illumina’s patterned flow cell, making it accessible to non-BGISEQ users. We believe that DocMF can provide valuable information for researchers from distinct communities.

## DATA AVAILABILITY

The data that support the findings of this study have been deposited in the CNSA (https://db.cngb.org/cnsa/) of CNGBdb with accession code CNP0000723.

## SUPPLEMENTARY DATA

Supplementary Data are available at NAR online.

## ACKNOWLEDGEMENT

We would like to thank the China National GeneBank and several scientists from BGI Research, including Liang Xiao, Yuanqiang Zou, Junhua Li and Yuxiang Li, for their friendly support and constructive suggestions.

