## Supplementary for "DNB-Based On-Chip Motif Finding (DocMF): a High-Throughput Method to Profile Different Types of Protein-DNA Interactions"

### Supplementary Materials & Methods

#### PAM Screen in vivo

The spacer of the random PAM plasmid library was randomly selected from the CRISPR array in VeCas9 CRISPR locus , and a random sequence of NNNNNNN was added at its 3' end or 5' end, and constructed into a pMD19 vector(Takara) by Gibson assembling(NEB). Finally, we constructed a 20-fold random PAM plasmid library (about 320,000 clones). VeCas9 CRISPR locus was cloned into the pACYC-184 vector by Gibson assembling, which was selected from the upstream and downstream 1000 bp of the CRISPR array and effector. The sequence was derived from the original strain genome. The screening experiment was performed by co-transforming the pACYC184 recombinant plasmid containing VeCas9 CRISPR locus and the 3'-end or 5'-end random PAM plasmid library in E. coli, and co-transforming a pACYC184 blank plasmid with a random PAM plasmid library as a negative control. Subsequently, the cloned plasmid was extracted, and the target PAM sequence was amplified, and the PAM recognition sequence was obtained by second-generation sequencing and bioinformation analysis.

| Name | Sequence |
|---|---|
| PAM_oligo_1 | TGTGAGCCAAGGAGTTGGCCTAGGCAATTGTCTTCTAAGACCGCTTGGCCTCCGACTT |
| PAM_oligo_2/1 | TTCGGTAGCAGTTCCCTTTTGAGNNNNNNNNNNNNNNNNAAGTCGGAGGCCAAGCGGTCT |
| PAM_oligo_2/2 | AGACCGCTTGGCCTCCGACTTNNNNNNNNNNNNNNNNCTCAAAAGGGAACTGCTACCGAA |
| PAM_oligo_3 | GAACGACATGGCTACGATCCGACTTNNNNNNNNNNNNNNNNTTCGGTAGCAGTTCCCTTT |
| Splint oligo | GCCATGTCGTTCTGTGAGCCAAGG |
| VeCas9-crRNA-F | TTCTAATACGACTCACTATAGGCTC |
| VeCas9-crRNA-R | GAAAAATTACAGAGTACTAAAACTTCGGTAG |
| VeCas9-TracrRNA-L-F | TTCTAATACGACTCACTATAGGAAAGG |
| VeCas9-TracrRNA-L-R | AACTACTTAAAAGTAGTGATTTCGACATAAAG |
| VeCas9-site-Cas9-F | TTCTAATACGACTCACTATAGGAAAAGGGAACTGCTACCGAAGTTTTAGAGCTAGA |
| Cas9-sgRNA-R | AAAAGCACCGACTCGGTGCCACT |
| VeCas9-TracrRNA-L | TTCTAATACGACTCACTATAGGAAAGGTTACAGAATCTACTAAAATAAGACTTTATGTCGAAATCACTACTTTTAAGTAGTT |
| VeCas9-crRNA | TTCTAATACGACTCACTATAGGCTCAAAAGGGAACTGCTACCGAAGTTTTAGTACTCTGTAATTTTTC |
| VeCas9-spacer-sgRNA-L | TTCTAATACGACTCACTATAGGCTCAAAAGGGAACTGCTACCGAAGTTTTAGTACTCTGTAATTTTTCaaaAAAGGTTACAGAATCTACTAAAATAAGACTTTATGTCGAAATCACTACTTTTAAGTAGTT |
| VeCas9-spacer-site-Cas9-sgRNA | TTCTAATACGACTCACTATAGGAAAAGGGAACTGCTACCGAAGTTTTAGAGCTAGAAATAGCAAGTTAAAATAAGGCTAGTCCGTTATCAACTTGAAAAAGTGGCACCGAGTCGGTGCTTTT |
| BvCpf1-crRNA-F | TTCTAATACGACTCACTATAGGAATTTCTACTATTGTAGATTTCGGTAGCAGTTCCCTTTTGAGC |
| BvCpf1-crRNA-R | GCTCAAAAGGGAACTGCTACCGAAATCTACAATAGTAGAAATTCCTATAGTGAGTCGTATTAGAA |

**Supplementary Table 1.** DNA oligo sequences used in this study

| Name | Ortholog | Complete sequence | Spacer |
| --- | --- | --- | --- |
| SpCas9 sgRNA | SpCas9 | AAAAGGGAACUGCUACCGAAGUUUUAGAGCUAGAAAUAGCAAGUUAAAAUAAGGCUAGUCCGUUAUCAACUUGAAAAAGUGGCACCGAGUCGGUGC | CTCAAAAGGGAACTGCTACCGAA |
| VeCas9 sgRNA-1 | VeCas9 | CUCAAAAGGGAACUGCUACCGAAGUUUUAGUACUCUGUAAUUUUUCaaaAAAGGUUACAGAAUCUACUAAAAUAAGACUUUAUGUCGAAAUCACUACUUUUAAGUAGUU | CTCAAAAGGGAACTGCTACCGAA |
| BvCpf1 crRNA | BvCpf1 | AAUUUCUACUAUUGUAGAUUUCGGUAGCAG UUCCCUUUUG AGC | CTCAAAAGGGAACTGCTACCGAA |
| dCas9 S. pyogenes sgRNA | dCas9 S. pyogenes | AAAAGGGAACUGCUACCGAAGUUUUAGAGCUAGAAAUAGCAAGUUAAAAUAAGGCUAGUCCGUUAUCAACUUGAAAAAGUGGCACCGAGUCGGUGC | CTCAAAAGGGAACTGCTACCGAA |

**Supplementary Table 2.** guide RNA and crRNA used in this study

| Name | Sequence |
|---|---|
| M13R300 | GTCAGTGAGCGAGGAAGC |
| M13F500 | TGCCACCTGACGTCTAAG |
| PAMveriR | TTCGGTAGCAGTTCCCTTTTGAGC |
| VeCas9-PAMveriF1 | GCTCAAAAGGGAACTGCTACCGAAGGTAGTCaATCTCTGGAAGATCCGCG |
| VeCas9-PAMveriF2 | GCTCAAAAGGGAACTGCTACCGAAGGTAAGAaATCTCTGGAAGATCCGCG |
| VeCas9-PAMveriF3 | GCTCAAAAGGGAACTGCTACCGAAGGTAGGAaATCTCTGGAAGATCCGCG |
| VeCas9-PAMveriF4 | GCTCAAAAGGGAACTGCTACCGAAACAGAACaATCTCTGGAAGATCCGCG |
| VeCas9-PAMveriF5 | GCTCAAAAGGGAACTGCTACCGAAACAGCATaATCTCTGGAAGATCCGCG |
| VeCas9-PAMveriF6 | GCTCAAAAGGGAACTGCTACCGAAACAAGCCaATCTCTGGAAGATCCGCG |
| VeCas9-PAMveriF7 | GCTCAAAAGGGAACTGCTACCGAACTGCAGCaATCTCTGGAAGATCCGCG |
| VeCas9-PAMveriF8 | GCTCAAAAGGGAACTGCTACCGAATGCAATTaATCTCTGGAAGATCCGCG |
| VeCas9-PAMveriF9 | GCTCAAAAGGGAACTGCTACCGAACCAATTGaATCTCTGGAAGATCCGCG |
| VeCas9-PAMveriF10 | GCTCAAAAGGGAACTGCTACCGAACCATTACaATCTCTGGAAGATCCGCG |
| VeCas9-PAMveriF11 | GCTCAAAAGGGAACTGCTACCGAAACGCATTaATCTCTGGAAGATCCGCG |
| VeCas9-PAMveriF12 | GCTCAAAAGGGAACTGCTACCGAAAGCGCGCaATCTCTGGAAGATCCGCG |
| VeCas9-PAMveriF13 | GCTCAAAAGGGAACTGCTACCGAAGGCATAAaATCTCTGGAAGATCCGCG |
| VeCas9-PAMveriF14 | GCTCAAAAGGGAACTGCTACCGAAGATCTAAaATCTCTGGAAGATCCGCG |
| VeCas9-PAMveriF15 | GCTCAAAAGGGAACTGCTACCGAAACTCCCAaATCTCTGGAAGATCCGCG |
| VeCas9-PAMveriF16 | GCTCAAAAGGGAACTGCTACCGAAAAATCATaATCTCTGGAAGATCCGCG |
| VeCas9-PAMveriF17 | GCTCAAAAGGGAACTGCTACCGAAAAACCGGaATCTCTGGAAGATCCGCG |
| VeCas9-PAMveriF18 | GCTCAAAAGGGAACTGCTACCGAAAAACCAAaATCTCTGGAAGATCCGCG |
| PAMveriF | ACGCCAAGTTTGCACGCCTGCCGTTCGACGATtGTAGTAGCTCAAAAGGGAACTGCTAC |
| VeCas9-PAMveriR1 | ACTCGGTACGCGCGGATCTTCCAGAGATtRKYYTRNTTCGGTAGCAGTTCCCTTTTGAG |
| VeCas9-PAMveriR2 | ACTCGGTACGCGCGGATCTTCCAGAGATtNNttTNNTTCGGTAGCAGTTCCCTTTTGAG |
| VeCas9-PAMveriR3 | ACTCGGTACGCGCGGATCTTCCAGAGATtNNctTNNTTCGGTAGCAGTTCCCTTTTGAG |
| VeCas9-PAMveriR4 | ACTCGGTACGCGCGGATCTTCCAGAGATtNNccTNNTTCGGTAGCAGTTCCCTTTTGAG |
| VeCas9-PAMveriR5 | ACTCGGTACGCGCGGATCTTCCAGAGATtNNtcTNNTTCGGTAGCAGTTCCCTTTTGAG |
| VeCas9-PAMveriR6 | ACTCGGTACGCGCGGATCTTCCAGAGATtNNYYgNNTTCGGTAGCAGTTCCCTTTTGAG |
| VeCas9-PAMveriR7 | ACTCGGTACGCGCGGATCTTCCAGAGATtNNYYaNNTTCGGTAGCAGTTCCCTTTTGAG |
| VeCas9-PAMveriR8 | ACTCGGTACGCGCGGATCTTCCAGAGATtNNYYcNNTTCGGTAGCAGTTCCCTTTTGAG |
| VeCas9-PAMveriR9 | ACTCGGTACGCGCGGATCTTCCAGAGATtNNTTTNNTTCGGTAGCAGTTCCCTTTTGAG |
| VeCas9-PAMveriR10 | ACTCGGTACGCGCGGATCTTCCAGAGATtNNAATNNTTCGGTAGCAGTTCCCTTTTGAG |

(continued)

(continued)

| | |
|---|---|
| BvCpf1-PAMveriF1 | GCTCAAAAGGGAACTGCTACCGAACAAAATTaATCTCTGGAAGATCCGCG |
| BvCpf1-PAMveriF2 | GCTCAAAAGGGAACTGCTACCGAAAAGAATAaATCTCTGGAAGATCCGCG |
| BvCpf1-PAMveriF3 | GCTCAAAAGGGAACTGCTACCGAAAGAAAGAaATCTCTGGAAGATCCGCG |
| BvCpf1-PAMveriF4 | GCTCAAAAGGGAACTGCTACCGAAAGGAAAAaATCTCTGGAAGATCCGCG |
| BvCpf1-PAMveriF5 | GCTCAAAAGGGAACTGCTACCGAACTTTATTaATCTCTGGAAGATCCGCG |
| BvCpf1-PAMveriF6 | GCTCAAAAGGGAACTGCTACCGAAACCTAAAaATCTCTGGAAGATCCGCG |
| BvCpf1-PAMveriF7 | GCTCAAAAGGGAACTGCTACCGAANRRNNNNaATCTCTGGAAGATCCGCG |
| BvCpf1-PAMveriF8 | GCTCAAAAGGGAACTGCTACCGAANRYNNNNaATCTCTGGAAGATCCGCG |
| BvCpf1-PAMveriF9 | GCTCAAAAGGGAACTGCTACCGAANAAANNNaATCTCTGGAAGATCCGCG |
| BvCpf1-PAMveriF10 | GCTCAAAAGGGAACTGCTACCGAANARANNNaATCTCTGGAAGATCCGCG |
| BvCpf1-PAMveriF11 | GCTCAAAAGGGAACTGCTACCGAANYRANNNaATCTCTGGAAGATCCGCG |
| BvCpf1-PAMveriF12 | GCTCAAAAGGGAACTGCTACCGAANYRBNNNaATCTCTGGAAGATCCGCG |
| BvCpf1-PAMveriF13 | GCTCAAAAGGGAACTGCTACCGAANRYANNNaATCTCTGGAAGATCCGCG |
| BvCpf1-PAMveriF14 | GCTCAAAAGGGAACTGCTACCGAANRYBNNNaATCTCTGGAAGATCCGCG |

**Supplementary Table 3.** DNA oligos used for in vitro PAM validation

| Name | Sequence |
|---|---|
| pMD19T-VeCas9- -3'PAM-R | CGCGGATCTTCCAGAGATtNNNNNNNTTCGGTAGCAGTTCCCTTTTGAGC |
| pMD19T- VeCas9-- 3'PAM-F | CCTGCCGTTCGACGATtGTAGTAGCTCAAAAGGGAACTGC |
| pMD19T-VeCas9-5’PAM-F | CGCGGATCTTCCAGAGATtTTCGGTAGCAGTTCCCTTTTGAGC |
| pMD19T-VeCas9-5’PAM-R | CCTGCCGTTCGACGATtNNNNNNNGTAGTAGCTCAAAAGGGAACTGC |
| VeCas9--1F | GAAGATCATCTTATTAATCAGATAAAATATTTCTGTACCTATTCATTTCACACACC |
| VeCas9- -1R | CAATTTAACTGTGATAAACTACCGCATTAAGCTACGCACCTGTAAACCTTG |

**Supplementary Table 4.** DNA oligos used for in vivo PAM validation

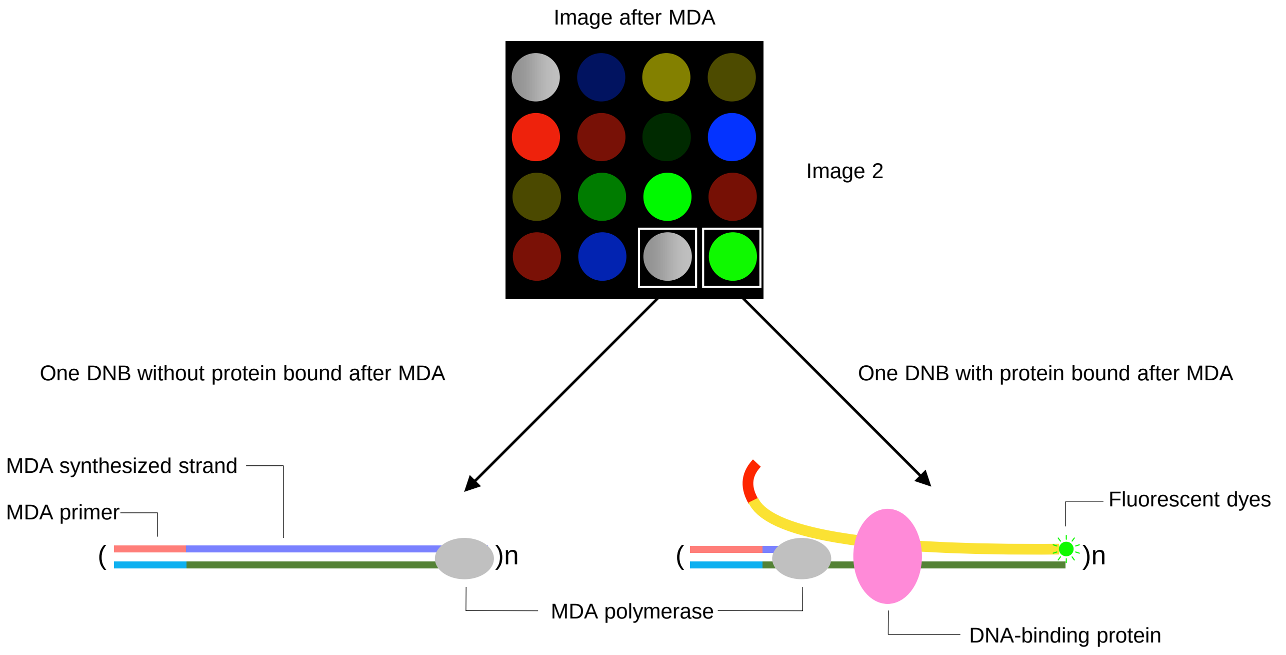


**Supplementary Figure 1.** Illustration of decreased fluorescence from a DNB after MDA in the presence of bound protein.

| Buffers | Standard | Formamide |
|---|---|---|
| Version | 0.5.0.10501 | 0.5.0.10501 |
| Reference | Ecoli | Ecoli |
| Cycle Number | 40 | 40 |
| Chip Productivity % | 88.88 | 85.08 |
| Avg Duplication Rate % | 96.72 | 94.9 |
| Q30 % | 92.83 | 90.23 |
| Lag % | 0.15 | 0.15 |
| Runon% | 0.15 | 0.12 |
| ESR % | 89.99 | 86.2 |
| Mapping Rate % | 99.3 | 99.27 |

**Supplementary Figure 2.** Sequencing performance before and after formamide treatment. Q30 refers to the percentage of basecall accuracy higher than 99.9%. Lag refers to delays in sequencing. RunOn refers to a sequencing process running faster than it is intended. These issues can be caused by incomplete polymerization or cleavage.

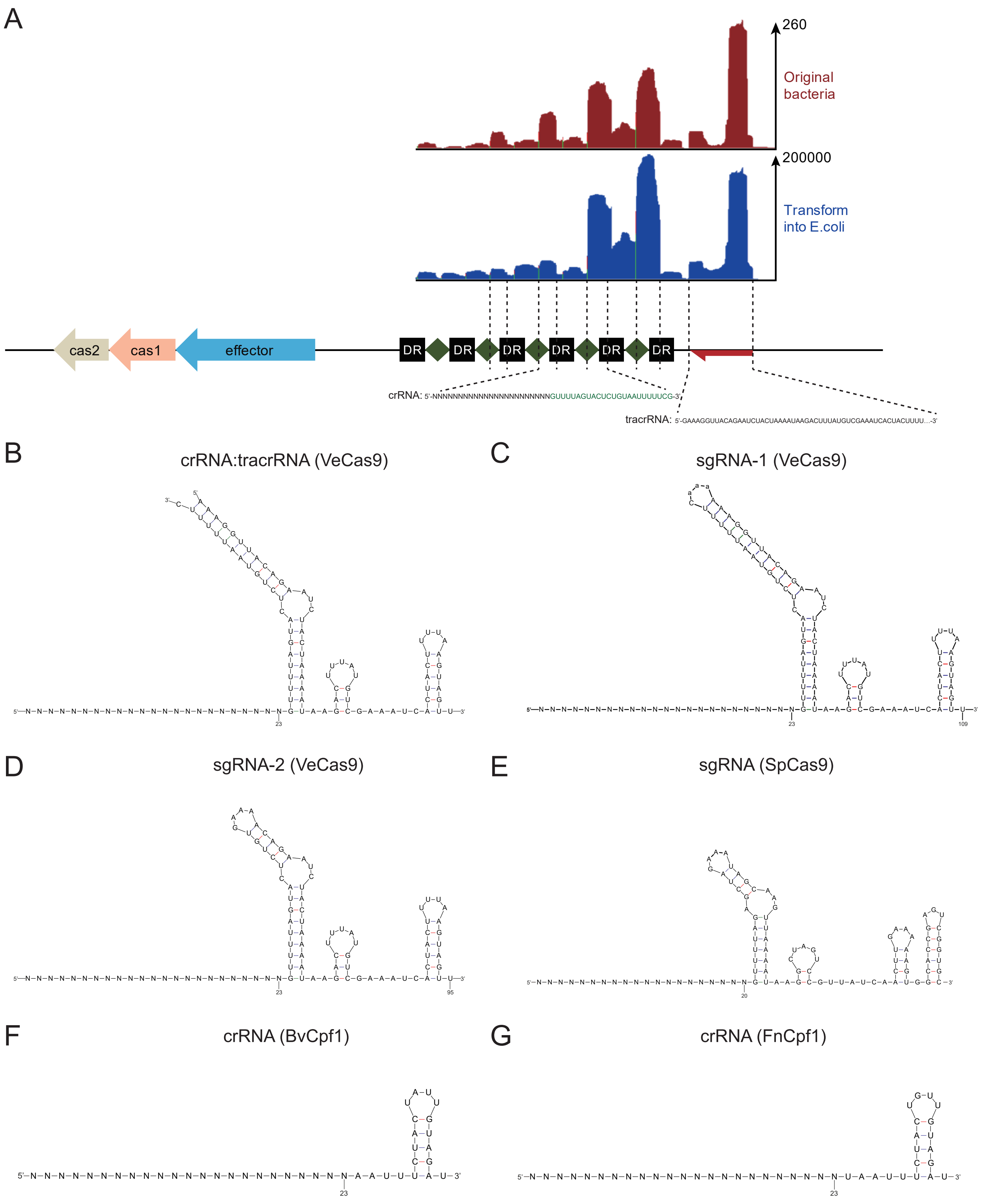


**Supplementary Figure 3.** (A) Small RNA-seq of VeCas9 in both original bacteria and E.coli containing all expression components. Both show robust expression of the CRISPR array. Additionally, the predicted tracrRNA close to the CRISPR array expresses well. (B) The co-folding prediction of crRNA and tracrRNA for VeCas9. (C) Folding prediction of sgRNA-1 for VeCas9. (D) Folding prediction of sgRNA-2 for VeCas9. (E) Folding prediction of common single guide RNA for SpCas9. (F) Folding prediction of crRNA for BvCpf1. (G) Folding prediction of crRNA for FnCpf1.

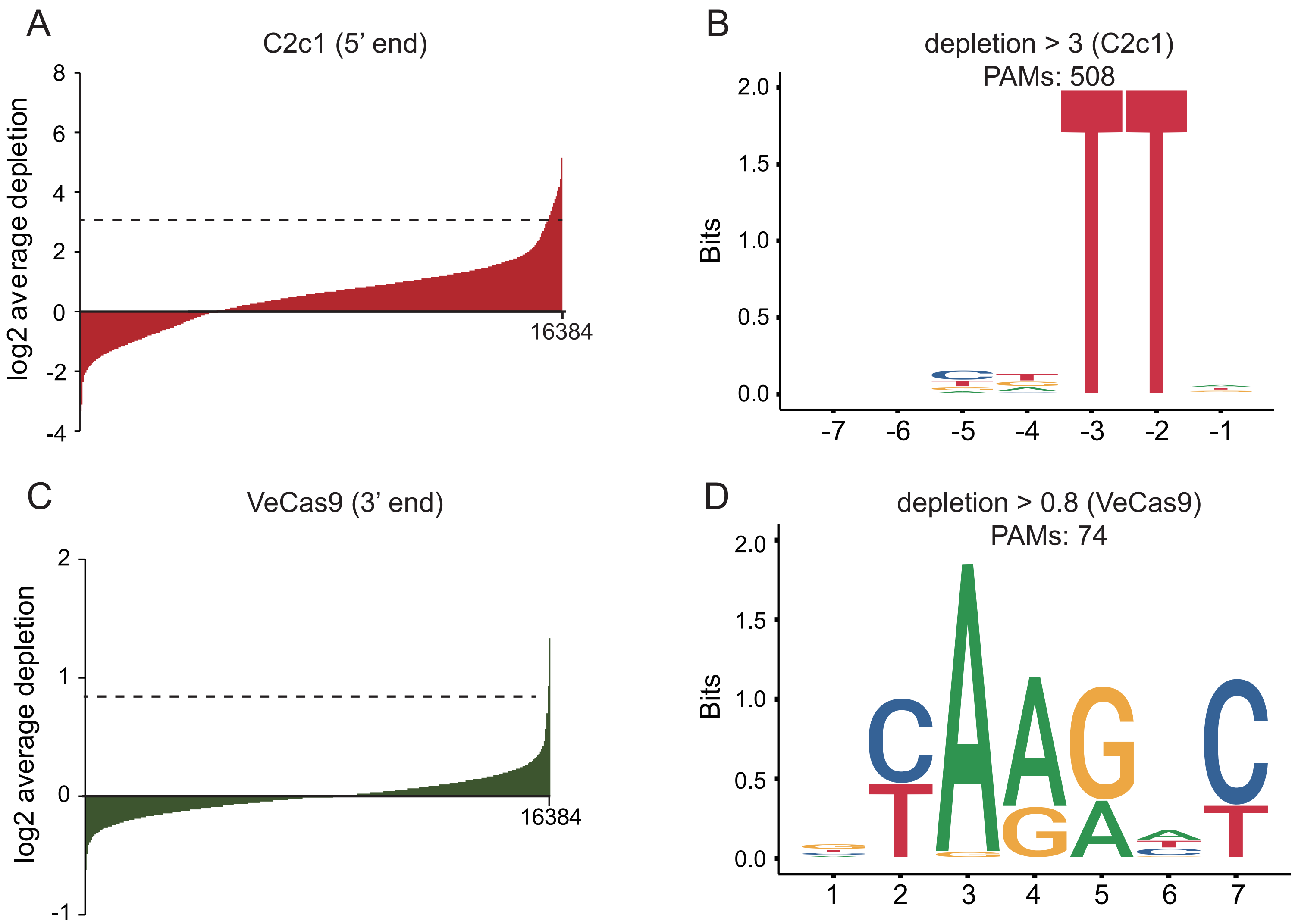


**Supplementary Figure 4.** (A) The sequence depletion from the 5'-end library for C2c1 reveals that sequences above a threshold of 3 significantly decline in this assay. Meanwhile, these differential sequences are pooled to generate PAM sequence. (B) Sequence logo for C2c1, which is generated by all sequences above the depletion threshold of 3, shows a TTN PAM. (C) The sequence depletion from the 3'-end library for VeCas9. There are a few sequences with considerable differences from others. (D) Sequence logo for VeCas9, produced by sequences above a depletion threshold of 0.8, shows a NYARRNY PAM.

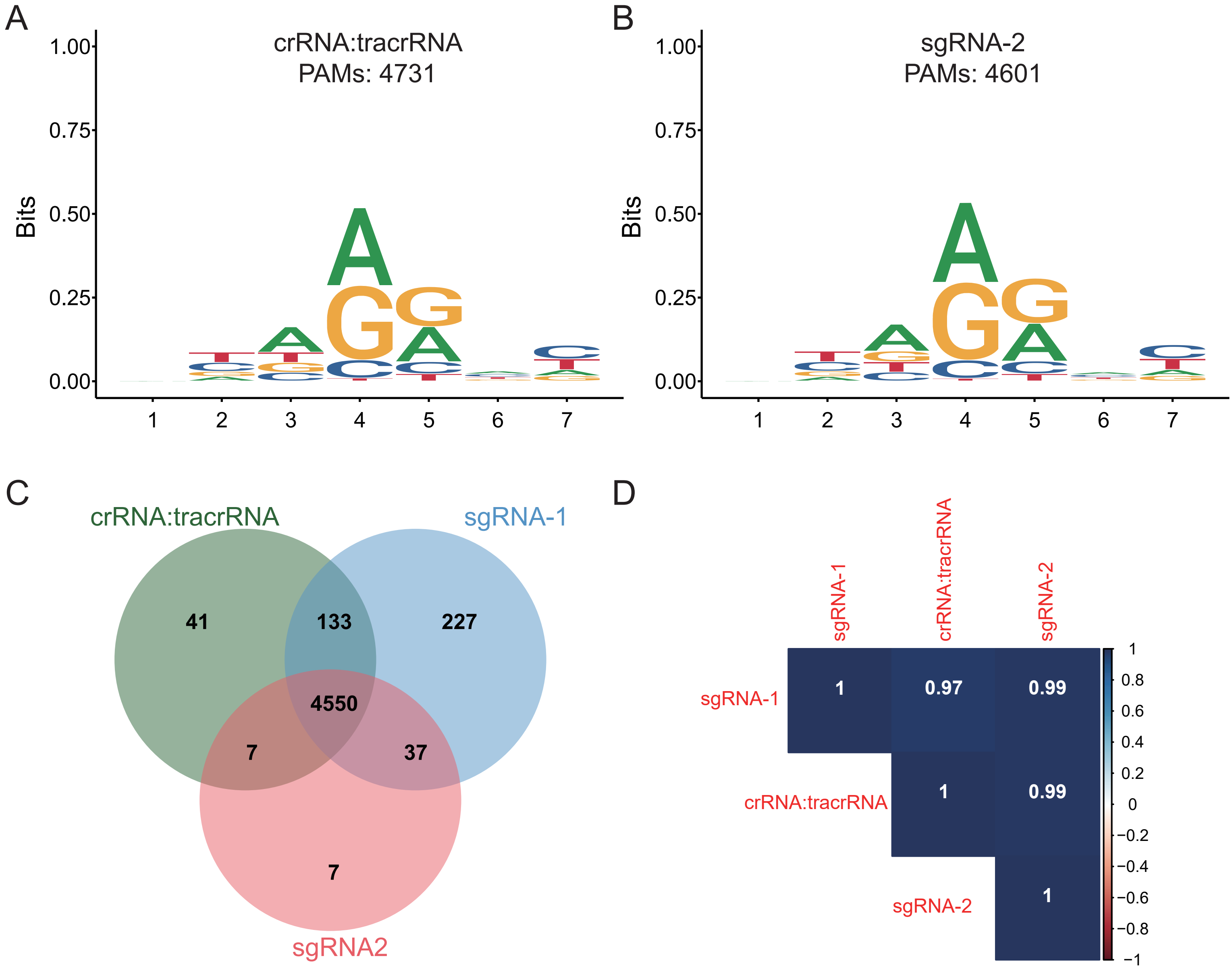


**Supplementary Figure 5.** (A) Sequence logo for VeCas9, which is detected by DocMF using the combination of crRNA and tracrRNA, shows an identical PAM with that using sgRNA-1. (B) Sequence logo for VeCas9 using sgRNA-2. (C) The Venn diagram of detected positive 7-nt sequences among three groups. Despite a slight difference among them, the vast majority are identical. (D) The heatmap of the Pearson correlation coefficient shows a strong correlation among three groups, which indicates DocMF platform reproducibility.

A

| Position 3 | Ratio | Position 4 | Ratio | Position 4 | | Position 5 | Ratio |
|---|---|---|---|---|---|---|---|
| A | 47% | A | 41% | A | 44% | A | 38% |
| | | T | 3% | | | T | 8% |
| | | C | 17% | | | C | 12% |
| | | G | 39% | | | G | 42% |
| T | 18% | A | 49% | T | 2% | A | 44% |
| | | T | 0.50% | | | T | 4% |
| | | C | 0.40% | | | C | 6% |
| | | G | 50% | | | G | 45% |
| C | 16% | A | 45% | C | 10% | A | 38% |
| | | T | 0.30% | | | T | 5% |
| | | C | 1% | | | C | 10% |
| | | G | 53% | | | G | 46% |
| G | 19% | A | 46% | G | 44% | A | 34% |
| | | T | 0.90% | | | T | 8% |
| | | C | 11% | | | C | 15% |
| | | G | 43% | | | G | 43% |

B

| Position -2 | Ratio | Position -3 | Ratio | Position -3 | | Position -4 | Ratio |
|---|---|---|---|---|---|---|---|
| A | 10% | A | 0 | A | 11% | A | 21% |
| | | T | 43% | | | T | 45% |
| | | C | 57% | | | C | 19% |
| | | G | 0 | | | G | 15% |
| T | 54% | A | 18% | T | 36% | A | 17% |
| | | T | 35% | | | T | 46% |
| | | C | 35% | | | C | 21% |
| | | G | 13% | | | G | 15% |
| C | 27% | A | 5% | C | 44% | A | 18% |
| | | T | 36% | | | T | 39% |
| | | C | 52% | | | C | 28% |
| | | G | 8% | | | G | 14% |
| G | 9% | A | 0% | G | 9% | A | 13% |
| | | T | 41% | | | T | 52% |
| | | C | 59% | | | C | 31% |
| | | G | 0% | | | G | 3% |

**Supplementary Figure 6.** (A) The base frequency for VeCas9 and the relevance of either position 3 and 4 or position 4 and 5. (B) The base frequency for BvCpf1 and the relevance of either position -2 and -3 or position -3 and -4.
